# Subretinal aspects of the optoretinographic response

**DOI:** 10.1101/2025.09.18.676889

**Authors:** Reddikumar Maddipatla, Maciej M. Bartuzel, Ewelina Pijewska, Christopher S. Langlo, Robert J. Zawadzki, Ravi S. Jonnal

## Abstract

Water movement in the living human retina and its regulation are important components of the tissue’s structural integrity, optical properties, and homeostasis. In the outer retina there is a continuous flow of water from the subretinal space, through the retinal pigmented epithelium, and into the choroid. This flow is disrupted acutely in disorders such as retinal detachment and central serous retinopathy, and is also known to reduce dramatically with age and age-related macular degeneration. Optoretiongraphy is an emerging technique for measuring neural function in the retina by monitoring nanometer-scale deformations of the membranes of photoreceptors. These deformations have been hypothetically attributed, in part, to osmotic shifts that cause water to move into and out of the photoreceptor outer segment after light stimulation. In the present work, we describe a method for measuring changes in the lengths of the cone outer segment and subretinal space in parallel and results showing that light stimuli change the volume of the subretinal space. These results are consistent with earlier *ex vivo* measurements of its light-induced hydration. The magnitude of the latter changes depend on the rate of water clearance from the subretinal space, and thus may serve as an indicator of the health of the water transport system. In addition, they may help us understand the mechanisms underlying the photoreceptor optoretinogram. These findings add to a growing understanding of the ways in which light exposure leads to transient reconfigurations of the outer retinal layers lasting milliseconds to hours.

## 2 Introduction

Aqueous humor (or simply aqueous) is a fluid secreted by the eye’s ciliary body, which lies in a ring on the inside of the eyeball, surrounding the lens. Its secretion and presence in the eye have several functions: transport of nutrients to the avascular cornea and maintenance of intraocular pressure and eye shape[1], which requires that the aqueous be drained as well. The main routes of drainage are through the trabecular meshwork and Schlemm’s canal, but a small fraction of it ( ≪ 1 %) moves through the vitreous body and into the retina via aquaporin-4 (AQP4) channels in the vitreous-facing end feet of Muller cells[2] and ultimately into the extracellular matrix (ECM) of the subretinal space (SRS). From the SRS, fluid is cleared primarily through active transport across the retinal pigmented epithelium (RPE) and pulled into the choriocapillaris by osmotic pressure. The full functional significance of this flow is unknown, but it has been shown that it is necessary for the adhesion of the neural retina to the underlying RPE[3].

Light-induced movement of water in the outer retina has been investigated in a number of animal models[4, 5, 6, 7, 8, 9] but it has not yet been measured in the living human eye. Accumulation of fluid in the retina is associated with a number of retinal disorders and diseases, such as retinal detachments, central serous retinopathy, age-related macular degeneration (AMD), Stargardt’s disease, and diabetic retinopathy[3, 10, 11].

Currently, in ophthalmic clinics, diseases affecting photoreceptors, such as AMD and geographic atrophy (GA), retinitis pigmentosa (RP), and ABCA4-associated cone-rod dystrophy, are assessed by observing structural abnormalities in the retinal layers using optical coherence tomography (OCT). However, assessment of retinal function may provide more timely information about the progression of these diseases, improving clinical management and streamlining drug discovery. Traditionally, photoreceptor function is assessed using visual acuity, visual field tests, and ERG. These methods are indispensable but they are not able to provide simultaneous structural and functional information. Optoretinography (ORG) is an emerging class of methods designed to measure stimulus-evoked changes in retina using noninvasive optical imaging. Photoreceptor structure and functional response have been measured together in cones [12, 13, 14, 15, 16] and rods [17, 18] using advanced ORG systems with adaptive optics (AO) or digital aberration correction (DAC). These approaches measure responses from single cells, most commonly by monitoring the phase difference between outer segment tips (COST or ROST) and the inner-outer segment junction (ISOS). However, clinical translation of AO-based imaging systems is hampered by their requirements of costly components, large amounts of space, multiple skilled operators, large capacities for data processing and storage, and lengthy measurement durations. Fortunately, novel, proto-clinical ORG approaches using custom OCT systems very similar to those currently employed in the clinics have been reported recently[19, 20].

In the present work, we used a proto-clinical, velocity-based approach[19, 21], to measure ORG responses inside and outside the photoreceptors in parallel, including boundaries of the outer segment (ISOS and COST) and the retinal pigmented epithelium (RPE). Specifically, we measured changes in thickness of three layers: ISOS-COST, COST-RPE, and ISOS-RPE. In ORG research, the ISOS-COST distance has been used as a marker of outer segment (OS) length. Changes in the OS optical path length (Δ*OPL*_*OS*_) have been the predominant source of signal in the ORG literature. The two other lengths we studied do not have neat anatomical counterparts. The COST-RPE length includes extracellular matrix (ECM), RPE apical processes (AP), and potentially phagosomes containing shed OS discs, while the ISOS-RPE length includes those components along with the OS. The first of these, the layer bounded by COST and RPE, has sometimes been called the supracone space[22, 23]. Some investigators use the term subretinal space (SRS) to denote the region containing ECM, AP, and phagosomes, while others use it interchangeably with ECM.

The use of the conflicting prefixes “sub-” and “supra-” to describe material that is outer to the retina and cone, respectively, is a historical accident. The former originates in 19th century clinical ophthalmic histology[24, 25] while the latter originates in neurobiological histology[26, 27], where the outer retina is traditionally positioned at the top of micrographs. In OCT-based anatomical studies, which position the outer retina at the bottom of the image, “subretinal space” is an intuitive name for material lying below the neural retina. Accordingly, “subcone space” may be more intuitive than “supracone space”. Fortunately, both can be abbreviated “SCS”, which is what we will use to designate it.

To our knowledge, the other length investigated in this study, the layer bounded by ISOS and RPE, has no accepted name. We will refer to it as the ciliary zone (CZ).

In four subjects without retinal disease, we observed the following after stimulus flashes: (1) the OS exhibits a brief (*<*20 ms) contraction followed by a longer elongation; (2) the SCS length exibits a brief (*<*20 ms) elongation followed by a longer contraction, complementary to the OS change but with smaller magnitude; and (3) the CZ exhibits a pattern of change similar to to the OS but smaller in magnitude.

Light exposure is known to increase the volume and water content of the ECM in the SRS[4, 28], but the methods used to establish that effect in animal models–concentration tracers and osmometry–are not suitable for human studies. Light-evoked increase in ECM hydration is a potentially significant process because it takes place in the context of a complex, highly regulated system of water movement from the SRS through the RPE and into the choroid. That system is disturbed in a number of diseases of the RPE-Bruch’s membrane (BrM) complex, most notably age-related macular degeneration[29]. It is very likely that the profound changes seen in BrM permeability in the aging and disease-affected eye would affect the dynamics of light-evoked SRS hydration. Changes in SCS and CZ cannot be unambiguously attributed to changes in the volume of a particular anatomical structure, because both spaces consist of heterogeneous material, both cellular (OS and AP) and acellular (ECM, phagosomes). However, because light-evoked SRS hydration is a well-known phenomenon, we believe it is likely to play a role in light-evoked changes in SCS and CZ, and this hypothesis is supported by the observations described below.

## 3 Methods

The OCT system has been described in detail in previous publications[19, 21]. In short, it consists of a 100 kHz swept-source laser (Axsun; *λ* = 1060 nm; Δ*λ* = 100 nm), two galvanometric scanners (Cambridge), a transmissive reference arm, a balanced photoreceiver (Thorlabs), and a digitizer (Alazar). A 90/10 fiber coupler sent 90% of the light to the reference arm and 10% to the eye. The back-scattered light from the eye was combined with the reference light using a 50/50 fiber coupler, and a retroreflector was translated in one dimension to adjust the reference arm length. The power of the illumination source, measured at the cornea, was 1.8 mW[19, 21]. The stimulus channel consisted of a 555 nm LED, selected because that wavelength stimulates the L and M cones equally. The right eyes of four healthy subjects were imaged, aged 29-50 years old. The research was conducted in compliance with the Declaration of Helsinki and with approval from the University of California Davis Institutional Review Board. Prior to imaging, the angles and interocular pressures of each subject were measured to assure low risk of acute open-angle glaucoma due to dilation. Then eyes were dilated using tropicamide 1%. Subjects were dark-adapted for five minutes prior to imaging. In each subject, five measurements, arranged in iso-eccentric arcs, were taken at each of five eccentricities: 2, 4, 6, 8, and 10 degrees. Each measurement consisted of a series of 80 B-scans at the same location collected at a rate of 400 Hz. B-scans consisted of 250 A-scans each, with a sampling interval of 3 µm. A circular 1.2 deg flash was delivered 40 ms after measurement initiated. The flash intensity was set to bleach about 65 % of L- and M-pigment. OCT signal processing was done using standard SS-OCT signal processing methods. Retinal layer velocities were computed using previously reported methods[19, 21], by analyzing three consecutive B-scans at a time. B-scans were aligned using amplitude-based cross-correlation and phase-based bulk motion correction. ISOS, COST, and RPE layers were segmented in a semi-automated way, and phase velocities were calculated at each layer in each lateral point in the segmented region. Relative velocities between the layers were calculated next, and then integrated laterally over the stimulated region to obtain the length change. OS and SCS elongation/contraction rates were calculated by subtracting the ISOS velocity from COST velocity, and the COST velocity from RPE velocity respectively, while CZ velocity was defined as the sum of OS and SCS velocities.

## 4 Results

Consistent patterns of response were found in the OS, SCS, and CZ in all subjects. After stimulus onset, the OS band contracted within the first 15 ms, and then elongated from 20-40 ms. The SCS exhibited elongation and contraction, respectively, during these periods. The CZ exhibited changes similar to the OS but with smaller magnitude. An example of these characteristic behaviors is shown in Fig. 1, which shows the OS, SCS, and CZ dynamics for Subject 1 from a single trial at one location 2 deg temporal to the fovea.

**Figure 1:**
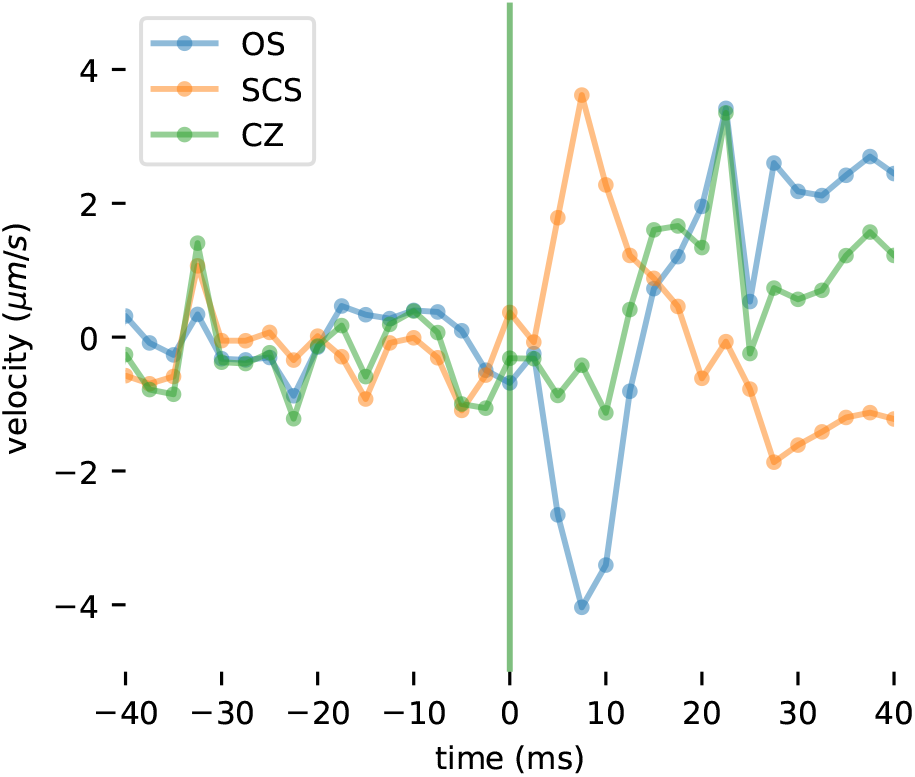
Changes in lengths of the OS (COST-ISOS), SCS (RPE-COST), and CZ (RPE-ISOS) in response to a visible stimulus. The relative velocity between ISOS and COST is the rate of contraction/elongation of the OS (blue line). As previously reported, after stimulus onset, the OS exhibits a rapid contraction followed by a slower period of elongation. The relative velocity of the SCS (orange line), between the COST and RPE, exhibits opposing changes: an initial elongation followed by a slower contraction. The CZ (green line) exhibits a change similar to the OS but with lower magnitude of contraction. Plotted data are the result of a single trial from Subject 1 at a location 2 deg temporal to the fovea. The vertical green line indicates stimulus onset.

Fig. 2 presents averaged responses from five different locations at 2^∘^ temporal from subject 9. Here, slight differences between the OS and SCS responses are visible. In this trial, the peak (contracting) velocity of the OS and the peak (expanding) velocity of the SCS both occurred 8 ms after stimulus onset, but in some trials the former preceded the latter by up to 2 ms. The magnitude of early OS contraction (∼− 4 µm s^−1^ in Fig. 2) was greater than the magnitude of early SCS expansion (∼ 3 µm s^−1^ in Fig. 2), and this was true in all subjects at all eccentricities. These differences suggest that though the OS and SCS responses have opposite signs, they are not fully complementary to each other.

**Figure 2:**
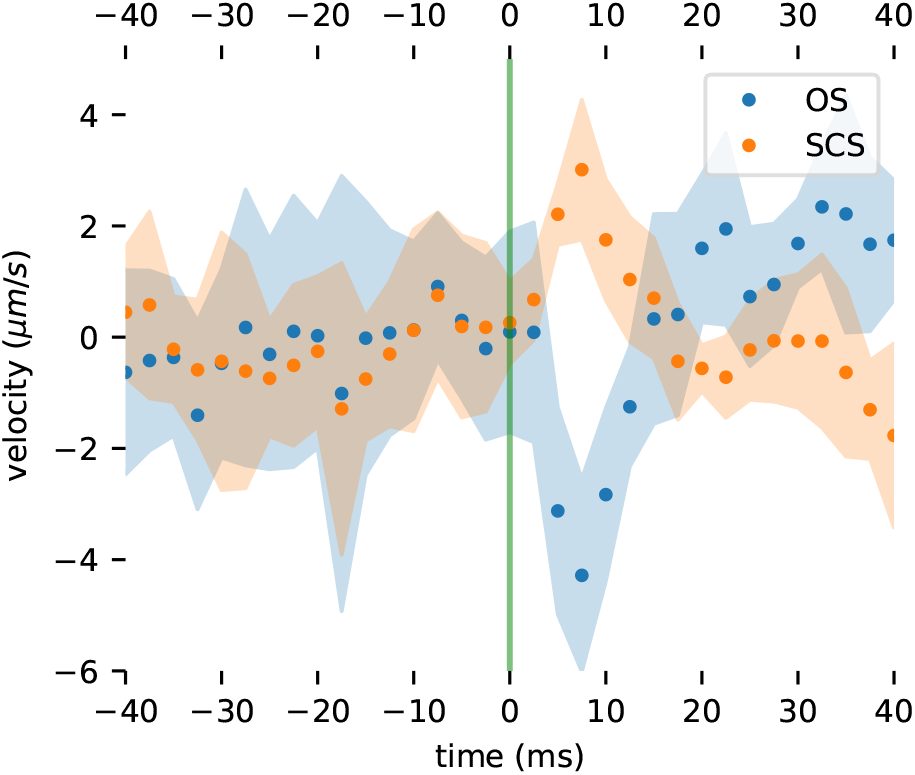
Averaging multiple trials to improve signal-to-noise ratio (SNR). Multiple trials were acquired at each eccentricity in each subject to facilitate the quantification of the OS and SCS changes. Averaging these together improves the SNR and visibility of the signals and their relationship. Plotted here are the average of five trials at 2T in Subject 9. Responses of the OS and SCS are shown (blue and orange circles, respectively) along with standard deviation of the trials (blue and orange shaded regions, respectively). While the two responses have opposite signs, they do not appear to be fully complementary. The initial SCS expansion is smaller in amplitude than the corresponding OS contraction.

Measurable functional responses of the OS and SCS were observed for all subjects at all eccentricities 2^∘^ to 10^∘^ (see Fig. 3). The absolute maximum velocity of OS and SCS varied among subjects, but tended to fall with increasing eccentricity.

**Figure 3:**
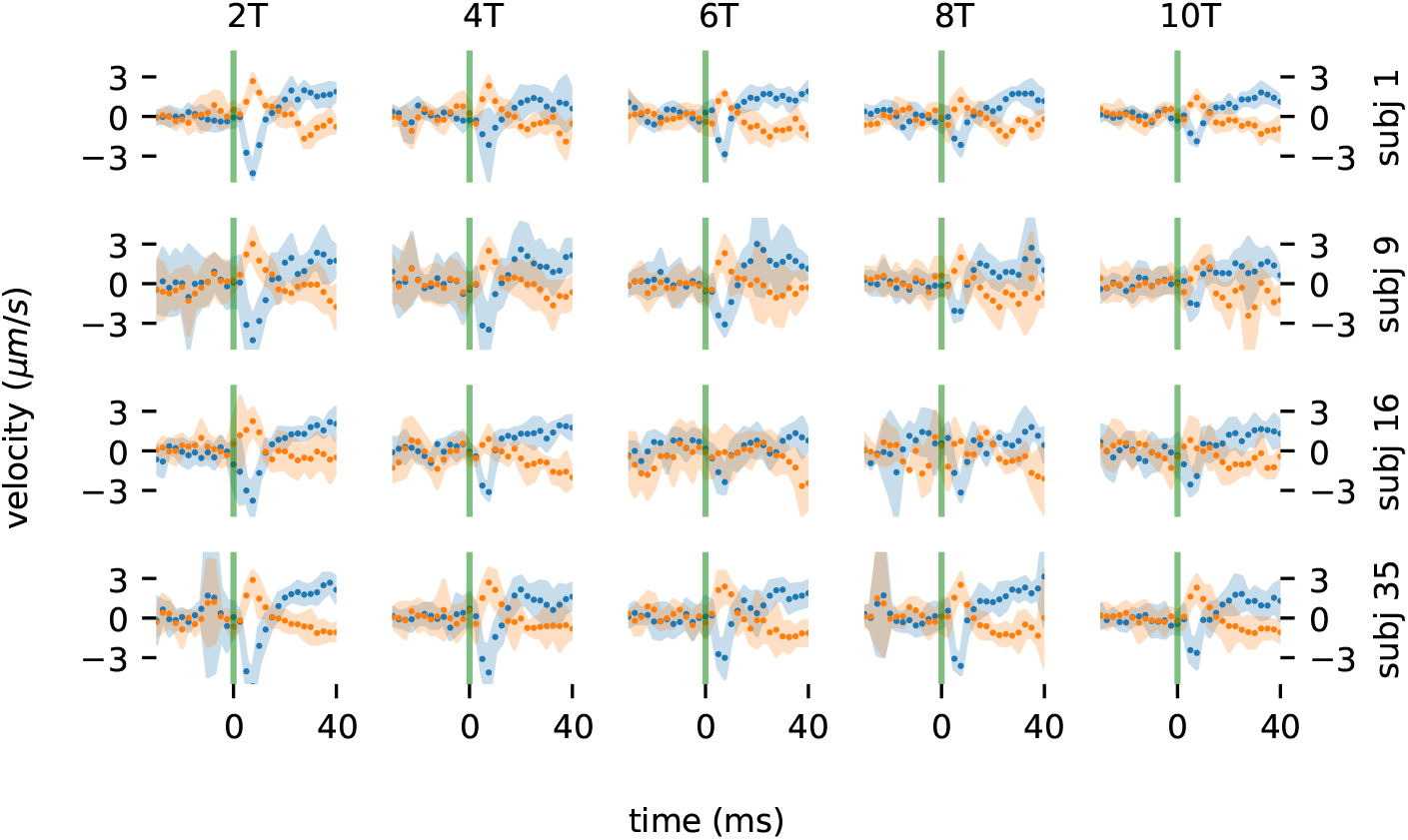
Responses from all subjects at all eccentricities. Plotted here are the rates of OS (blue) and SCS (orange) length change for all tested subjects and eccentricities. The individual responses and eccentricity-dependence of responses were consistent among subjects.

Responses were summarized by identifying extrema between 0 ms to 20 ms and computing the mean velocity between 20 ms to 40 ms. Identification of extrema (maximum or minimum velocity) was motivated by the observation that early responses had a visible trough (OS andCZ) or peak (SCS). Resulting summaries are shown in Fig. 4, with parameters determined at the same eccentricity offset horizontally for visibility.

**Figure 4:**
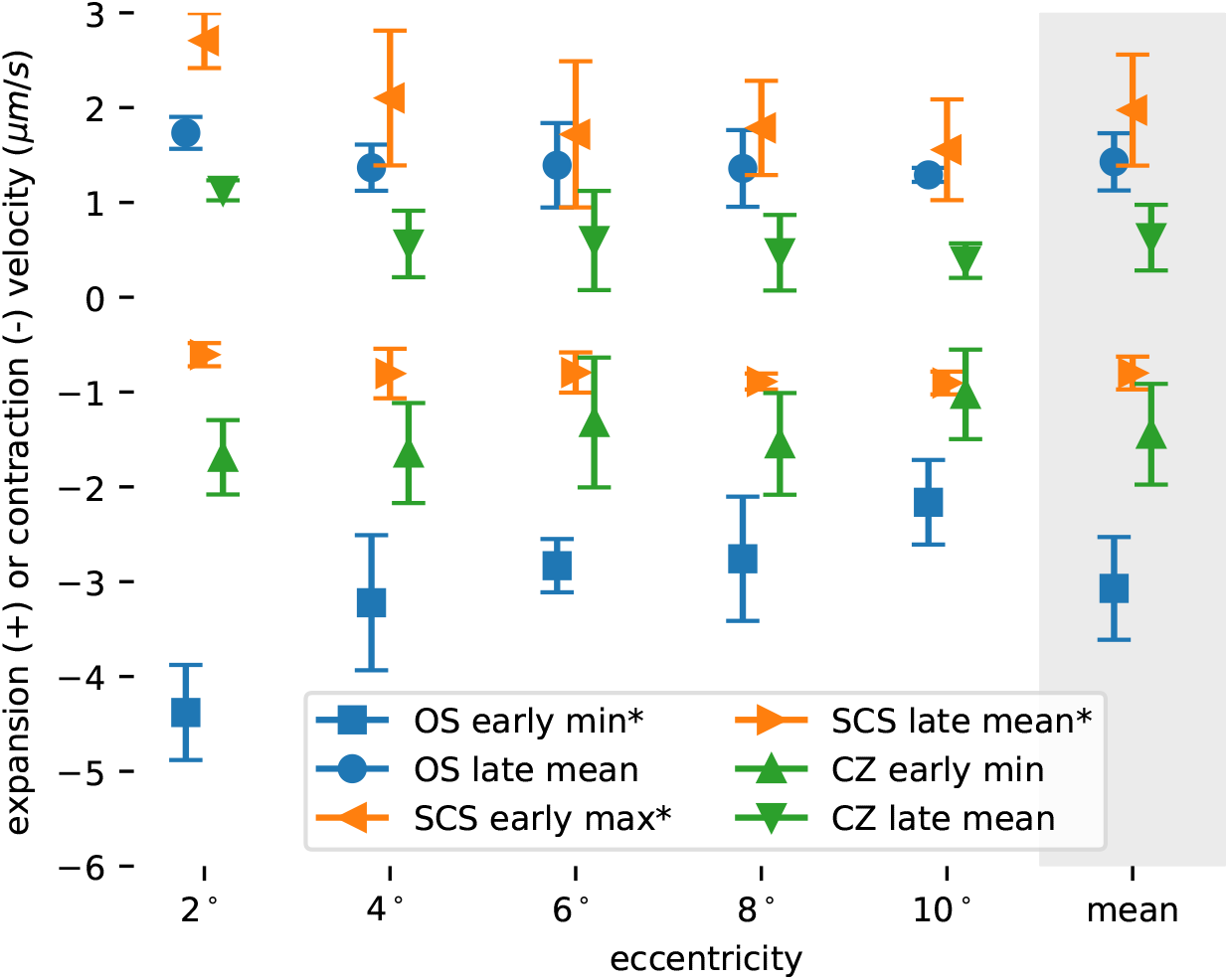
Response parameters as a function of eccentricity. Plotted here are summaries of the responses, averaged across subjects and plotted against eccentricity. Each response was quantified in the early (*<*20 ms) and late (20 ms to 40 ms) stages of response, using extrema or mean value, respectively. Each marker represents the average parameter across subjects, with error bars depicting the standard deviation. Three of the parameters (OS early min, SCS early max, and SCS late mean) exhibited statistically significant (*p <* 0.05) correlations with retinal eccentricity.

Error bars indicate the standard deviation among subjects. Of the six parameters computed three exhibited significant correlation with eccentricity: OS early minimum (*p* = 0.018), SCS early maximum (*p* = 0.032), and SCS late mean (*p* = 0.034).

To assess the reciprocity between changes in the OS and SCS, we first integrated the relative length changes of the OS and SCS and then added them together. If the two changes were reciprocal, then the sum of the two would remain near zero. Instead, we found that the sum, which corresponds to the ISOS to RPE distance, decreased over the first ∼20 ms and then increased thereafter. The pattern of changes resembled that of the OS itself, but with lower magnitude. To further reduce noise and verify that this effect was real, we averaged together the integrated and summed responses from 2^∘^ and 4^∘^ in each subject. The resulting responses from all four subjects are plotted in Fig. 5, and clearly show a lack of reciprocity and overall contraction and then expansion of the distance between ISOS and RPE.

**Figure 5:**
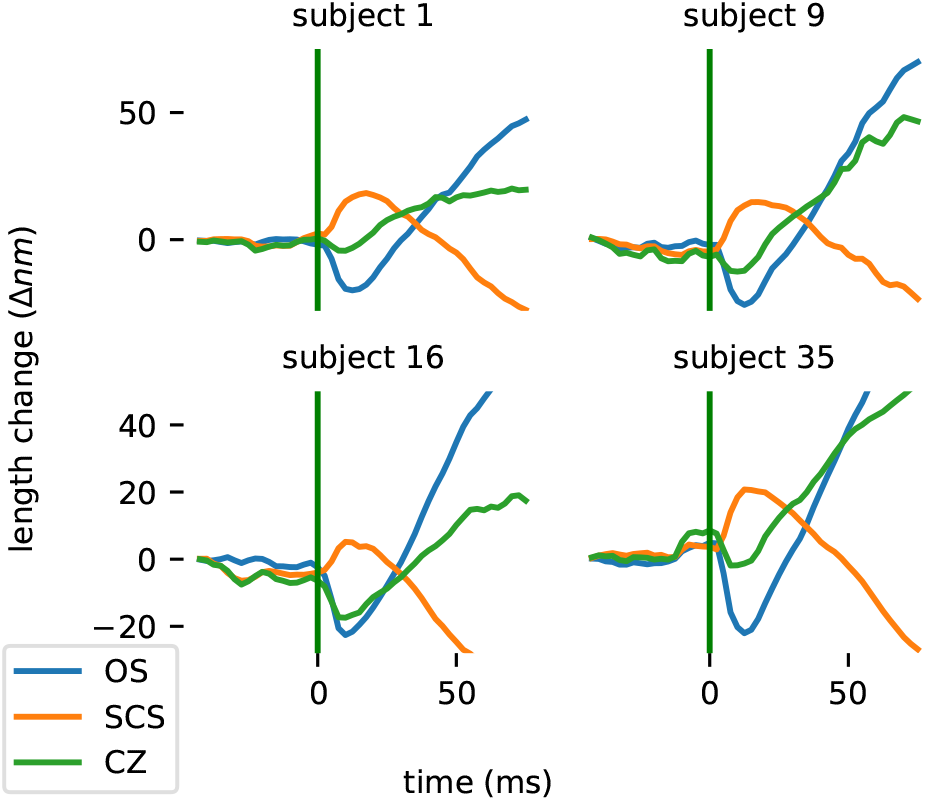
Lack of reciprocity of OS and SCS responses. Shown in these plots are the cumulative sums (numerical integrals) of relative velocities, which should correspond to length changes of the relevant zones. Length changes are plotted for the OS and the SCS, as well as the sum of these two responses, averaged across the 2 and 4 deg eccentricities, for each of the four subjects. In all subjects, the sum of the two individual zone responses is non-zero, indicating that the OS and SCS responses are not reciprocal.

## 5 Discussion

### 5.1 Summary of results

In the results above, six distinct aspects of the response were apparent. Hypothetical attribution of the early OS contraction and later OS elongation to separate mechanisms[16] suggests that the early ( ≤20 ms) and late (≥ 20 ms) stages of all the responses should be treated separately. In previous work, we quantified these stages using the maximum absolute velocity of the early stage and the mean velocity between 20 ms to 40 ms, and we follow the same approach here. The ranges below are due to variation among subjects and eccentricity, and the averages are across both. These six aspects are illustrated in Fig. 4.

#### 1. Early (≤20 ms) effects of light stimulus

a. Contraction of the OS with a most negative velocity ranged from −1.6 µm s^−1^ to −5.2 µm s^−1^, with an average of −3.1 µm s^−1^.
b. Elongation of the SCS with a maximum velocity ranged from 0.5 µm s^−1^ to 3.0 µm s^−1^, with an average of 2.0 µm s^−1^.
c. Contraction of the CZ with a most negative velocity ranged from −0.4 µm s^−1^ to −2.3 µm s^−1^, with an average of −1.4 µm s^−1^.

#### 2. Late (≥20 ms) effects of light stimulus

a. Elongation of the OS with a velocity ranged from 0.7 µm s^−1^ to 2.0 µm s^−1^, with an average of 1.4 µm s^−1^.
b. Contraction of the SCS with a velocity ranged from −0.4 µm s^−1^ to −1.2 µm s^−1^, with an average of −0.8 µm s^−1^.
c. Elongation of the CZ with a velocity ranged from 0.0 µm s^−1^ to 1.3 µm s^−1^, with an average of 0.6 µm s^−1^.

The observations of early OS contraction (1a) and late OS elongation (2a) corroborate similar findings from numerous groups[13, 15, 16, 19], as shown in Fig. 6 below. The remaining results above include three key findings: 1) the space between COST and RPE (SCS) exhibits a light-evoked change in thickness consisting of a rapid ∼20 ms expansion followed by a slower contraction; 2) the space between the ISOS and RPE (CZ) exhibits changes more like the OS itself, consisting of a rapid contraction followed by elongation; and 3) both components of the latter are smaller in magnitude than those of the OS change. All of these are visualized in Fig. 5. To our knowledge, this is the first report of light-evoked changes in SCS and CZ directly measured in the living human eye, though as we discuss below, the results do not conflict with previous findings by other groups. Changes in the length of CZ and SCS may involve the movement of water around the ECM or between the ECM and the cells that enclose it. Thus it will be useful to review some of what is known about water movement in the outer retina.

**Figure 6:**
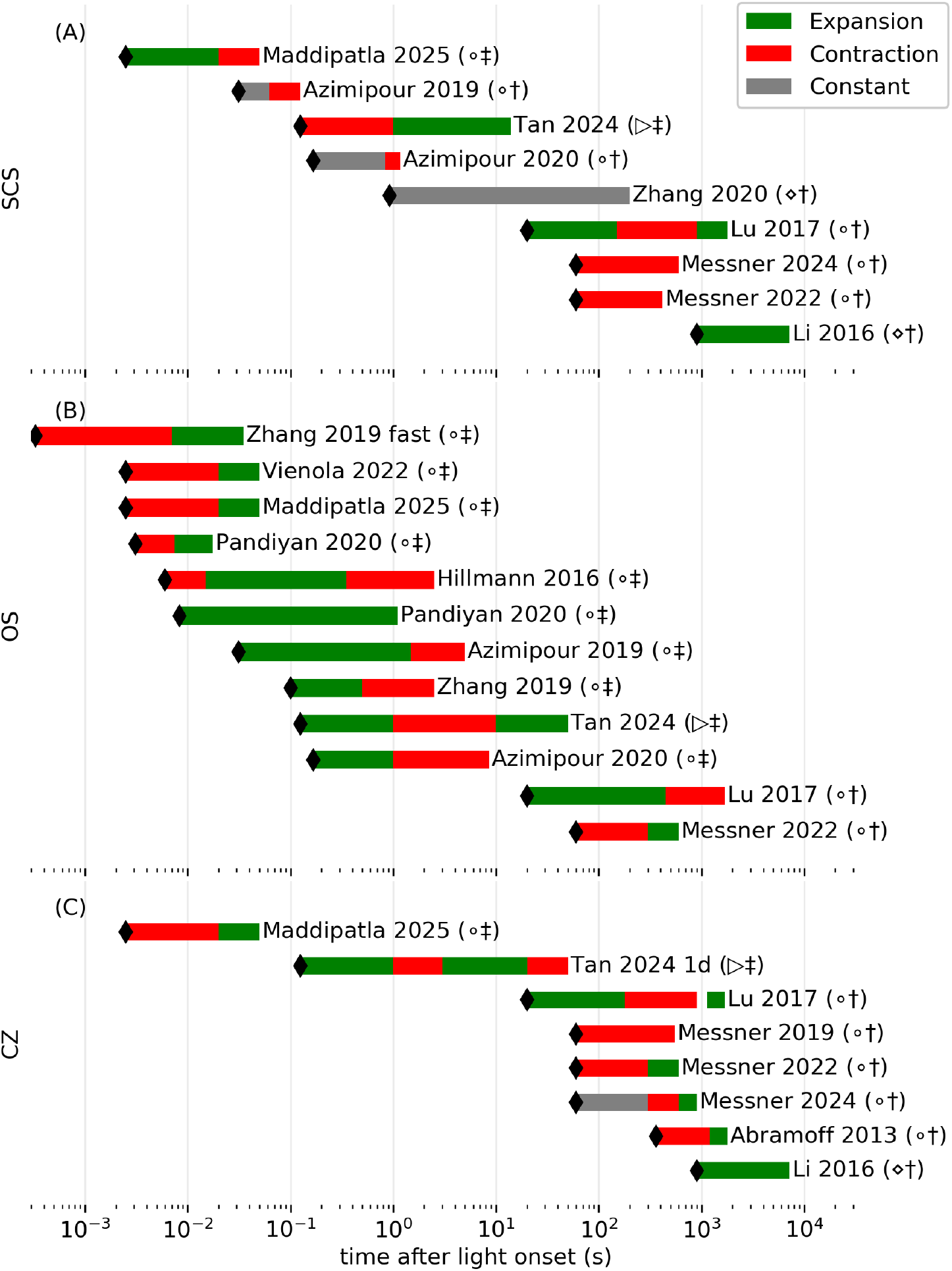
Review of light-evoked changes in length of outer retinal compartments. In the past decade, a number of investigators have used OCT to measure light-evoked changes in the thickness of various outer retinal structures. Black markers indicate the measurement bandwidth, and thus the earliest possible observation of elongation or contraction. Bars indicate the measurement duration and are colored green to indicate expansion of the structure, red to indicate contraction, and gray when no change was observed. ∘: human; ▹: rat;⋄: mouse; †: measurements using localization of bands in OCT amplitude; ‡: measurements using differences in OCT phase. See section 5.3 for detail and full citations.

### 5.2 Water movement in the SRS and light-evoked hydration of ECM

Light-evoked hydration of the SRS has been reported in frog[4] and chick[28, 6]. The magnitude this hydration, under bright stimuli, has been estimated to be 7 % to 17 %. The lower end of that range, 7 % is taken directly from changes in the concentration of the impermeant ECM tracer tetramethylammonium (TMA+)[28], whereas the upper end of that range is derived from a model devised by the same authors to correct for the effect of TMA+ diffusion[6].

While the initial rate of hydration was not reported, Fig. 3A in Li et al. (1994) suggests that it is ∼1.0 % s^−1^[28], or ∼2.5 % s^−1^ using the corrected hydration percentage of 17 %. We will proceed with the corrected value. In the locations studied in the present work, the distance between ISOS and RPE ranged from 30 µm to 50 µm. Using electron microscopy (EM) and semi-automated segmentation, investigators have shown that the 38% of the SRS consists of extracellular matrix, with OS and AP occupying 28% and 34%, respectively[30]. Assuming an ISOS-RPE length of 40 µm, and a 38% ECM composition, a 2.5 % s^−1^ increase in volume corresponds to an elongation rate of 380 nm s^−1^.

The elongation of CZ, averaged over subjects and eccentricities, was 600 nm s^−1^ (see item 2c above, as well as Fig. 4). The velocity derived from the 2.5 % s^−1^ rate of hydration, 380 nm s^−1^, is lower than this average value and significantly lower than the highest elongation velocities observed in subjects 9 and 35.

Several factors, potentially in concert, could underlie this shortfall:

1. The slab of tissue we have designated CZ includes the OS, which mostly elongates after stimuli. In rods the entirety of the OS is isolated from the ECM by membrane, and in cones a portion of the OS is isolated from the ECM. As with cones, rods have been shown to predominantly elongate after light stimuli[17]. If we assume a constant OS diameter, this elongation must entail an increase in the volume of the OS and thus an increase in its contribution to CZ volume. Presuming that the extracellular concentration of TMA+ ([TMA+]_*o*_) used to measure SRS hydration measured changes only in the ECM hydration, they would tend to underestimate the total expansion of the CZ. Our observation that the SCS contracts while the OS elongates supports this hypothesis, since it suggests that water is displaced from the SCS and into the the ECM between OSs, resulting in an increase in CZ volume that has no correlated reduction in the concentration of solutes (including tracers) in the ECM.
2. The stimulus dose used in the chick experiment may have been lower than what we used. The radiant exposures were 0.6 mJ*/*cm^2^ during the initial 10 s of stimulation, during which the monotonic 2.5 % s^−1^ hydration rate was observed in the chick experiment, whereas our radiant exposure was 0.98 mJ*/*cm^2^. This is an admitted approximation, and glosses over the spectral sensitivities of the chick’s five photopigments and our four photopigments as well as their use of a broad-spectrum halogen bulb and our use of a narrow 555 nm LED.
3. The average rate of hydration over seconds is 2.5 % s^−1^ but could be higher in the first 50 ms, and the deceleration of hydration rate may have been undersampled or not reported[28]. Indeed, a later linear systems model of SRS hydration suggested that diffusion of the tracer effectively convolves the hydration signal with a diffusion function, which would lead to an underestimate of the hydration rate[6].
4. The EM-based estimate of SRS fraction (38%) was derived from a single sample from a parafoveal location 3^∘^ to 4^∘^ in a single subject. However, eccentricity-dependent variation the comparative footprints of cone IS and cone OS implies that this percentage may vary geographically, with cone OS packed tightly in the fovea and more sparsely elsewhere and rod OS making up part of the difference.
5. Interspecific[31] or age-related[29] differences in RPE water transport could partly underlie the high CZ velocity we observed.

The 600 nm s^−1^ CZ elongation rate we observed corresponds to a volumetric flux of 1.13 µL*/*mm^2^*/*h, ∼10*×* higher than the maximum clearance flux measured in humans of 0.11 µL*/*mm^2^*/*h[32], which explains why the CZ swells in spite of the excess capacity of the RPE’s water transport system in normal conditions[3].

The origin of the water that causes the light-dependent CZ swelling reported here and earlier reports of light-dependent hydration[4, 28, 6] is uncertain. There is a continuous flow of water from the inner retina toward the outer retina[3]. Mechanisms of this transport include facilitated diffusion, paracellular flow, and osmotic/oncotic mechanisms[33]. The maximum volumetric flux from SRS, through RPE, and into the choriocapillaris has been estimated to be 0.12 µL*/*mm^2^*/*h to 0.31 µL*/*mm^2^*/*h in rabbits[3], 0.064 µL*/*mm^2^*/*h in dogs[34], 0.073 µL*/*mm^2^*/*h in cynomolgus monkeys[35], and 0.11 µL*/*mm^2^*/*h in humans[32]. It is important to note that normally, the rate of water transport is limited by the availability of water in the SRS, while the measurements above were taken when an abundance of fluid was available on the apical side of the the RPE, mirroring the situation obtained in serous retinopathy and some retinal detachments[3].

The influx of water into the SRS is determined in part by intraocular pressure (IOP)[3], which pushes water through the retina, possibly via AQP4 channels of Muller cells[2]. IOP, presumably, is unaffected by light stimuli. However, it is possible that the shifting concentrations of osmolytes in the SRS increases osmotic draw of water through Muller cells or paracellular routes. As mentioned above, elongation rate of 600 nm s^−1^ corresponds to a volumetric flux of 1.13 µL*/*mm^2^*/*h. It is possible that the outward route from the retina into the SRS could accommodate this flux, though the 10 *×* smaller maximum clearance rates of the RPE may suggest otherwise. If that route cannot supply the required flux, we must infer that the water moves from the RPE into the SRS. Such a reversal violates the conventional wisdom that outward water flow is required for retinal-RPE adhesion[3]. On the other hand, perhaps transient disruptions of this flow are tolerable, as experimental subretinal edema is quickly cleared in rabbits[36, 37] and central serous retinopathy is usually self-limiting and self-resolving[38]. Moreover, disabling the inward water transport system in the apical RPE appears to attenuate this hydration effect[4, 5].

All of the analysis above presumes that the sources of the OCT signal are boundaries of compartments that expand or contract with changes in hydration. If, on the other hand, these scatterers are free to move within a compartment and do so in response to light stimuli, interpretation as layer expansion and contraction may be incorrect. Melanosomes in the RPE are thought to be the source of contrast in the RPE[39], and thus our SCS and CZ measurements likely depend on the their location in the RPE cell. Melanosomes are thought to congregate near the apical surface of the RPE cell[40], but have also been observed within its apical processes[41, 42]. Light-driven translocation of melanosomes in amphibians and fish is widely reported, and has been observed in frogs using OCT[43]. Though mammals are not thought to have comparable retinomotor systems, passive movement of melanosomes within the RPE, perhaps associated with water movement, must be considered as a possible alternative explanation. Indeed, changes in light level have been shown to cause slow movement of melanosomes within the RPE cell in mice[42].

### 5.3 Synthesis of results with prior work

In the past fifteen years, many investigators have used OCT to study light-evoked changes in the outer retina. Much of this work is summarized in Fig. 6. Panels (A), (B), and (C) depict summaries of findings from the SCS, OS, and CZ, respectively. The abscissa represents time in log scale, and each row of each plot represents a single study. Each bar represents the time over which layer thickness changes were investigated, with green indicating expansion and red indicating compression for the relevant layer. We chose to depict only the direction of change (expansion/contraction) and not its magnitude, because of the presumed large variation in magnitudes among species and stimulus characteristics. In several of the rows of the figure, the specific layer depicted was not explicitly reported in the study but could be inferred from other results; our methodology, with reference to the relevant figures, will be described in passing. Most of these studies have focused on photoreceptor responses, mainly by monitoring the length of the outer segment as it is exposed to light stimuli 6 (B). These are presented for completeness but will not be discussed extensively here; much of the work has been reviewed elsewhere[44]. A few, however, have reported corresponding changes outside the photoreceptor including the layers investigated in this work, and we will describe that work in more detail here.

We have previously reported rapid changes in the amplitude of the OCT signal after light stimuli that are consistent with contraction of SCS around 0.1 s (Azimipour 2019 *†*)[14] and 1.0 s (Azimipour 2020 *†*)[17]. These consisted of apparent movement of an aspect of the RPE band inward, toward the ISOS, after stimuli. They are shown in Fig. 6 (A).

Zhang et al. (2020) used the OCT amplitude image along with Gaussian mixture modeling of the axial amplitude profile to discriminate ROST and RPE, and monitored the location of each with respect to BrM during light exposure. The difference between their locations corresponds to our SCS, albeit beneath rods, which are much more closely apposed to RPE. Interestingly, they showed that the thickness of SCS did not change after stimulation, but remained a constant ∼3.3 µm. This is shown in Fig. 6 (A).

Recently Tan et al. (2024) reported light-evoked changes in the rat SRS[8] on time scales of hundreds of milliseconds, using OCT phase. They defined the SRS thickness as the distance between the ELM and the RPE. In the OCT image of the rod-dominant rat retina, contributions from the ROST and RPE cannot be visually distinguished. However, an unsupervised learning method was employed to differentiate these based on light-evoked phase changes. Some of the pixels in the ROST-RPE complex were shown to exhibit movement, relative to the IS/OS, similar to what has been shown in human rods and cones using AO-OCT: an OS elongation occurring over hundreds of milliseconds followed by a contraction lasting seconds and another elongation lasting tens of seconds. These are depicted in Fig. 6 (B). Other pixels, thought to originate from the RPE, exhibit slower, monotonic movement away from IS/OS over tens of seconds. The investigators attributed the latter to slow, light-evoked expansion of the subretinal space. In this way they were able to monitor movements of the ISOS, ROST, and RPE relative to the ELM. By digitizing and subtracting responses of ISOS from ROST and ROST from RPE, we inferred the behaviors of CZ and SCS, respectively. SCS appeared to contract between 0.125 s to 1 s and elongate between 1 s to 14 s, as shown in Fig. 6 (A). CZ underwent a four-phase deformation, elongating between 0.125 s to 1 s, contracting between 1 s to 3 s, elongating again between 3 s to 20 s, and finally contracting between 20 s to 50 s. This response is depicted in Fig. 6 (C).

Other investigators have measured light-evoked changes outside of the photoreceptors over longer time scales. Abramoff et al. (2013) used the term outer segment equivalent length (OSEL) to describe what we here have called CZ[45]. They showed that CZ contracts between 6 min to 20 min and expands between 20 min to 30 min after light onset.

Li et al. (2016) showed differences between dark- and light-adapted mouse retina[46]. They reported an overall light-dependent swelling of the outer retina, bounded by ELM and the choroid, between 15 min to 120 min after light onset. From data shown in the paper, it is evident that the IS length does not change, which implies that the CZ experiences light-dependent swelling. Although not reported by them, it is evident from comparison of the dark- and light-adapted axial profiles (their Fig. 1E) that a portion of that expansion is seen between the ROST and RPE, consistent with light-dependent expansion of the SCS. These effects are shown in Fig. 6 (A) and (C).

Lu et al. measured such effects in humans between 20 s and 30 min following a bright stimulus flash that bleached varying amounts of rod and cone photopigment. Numbers below refer to trials in which 96 % of rhodopsin was bleached. In cone OS they observed an immediate elongation of 500 nm to 750 nm, followed by a recovery to baseline within a few minutes. Rod OS (ISOS-ROST distance) was found to elongate 400 nm to 800 nm over the first 6 min following the bleach, and return to baseline over the next 20 min. They reported parallel changes in the distance of RPE from ISOS during recovery from the bleach. This distance, which corresponds to our CZ, exhibited a triphasic response, elongating between 0.33 min to 3 min, contracting between 3 min to 15 min, and elongating again to baseline between 15 min to 30 min. From their measurements of rod OS and CZ we were able to digitize and estimate the change in SCS to consist of elongation between 0.33 min to 2.5 min, contraction between 2.5 min to 15 min, and elongation to baseline between 15 min to 30 min.

Three studies were conducted by Messner et al. using the amplitude image of a commercial OCT system. First, they measured light-evoked contraction of the ISOS-RPE distance, what we have referred to as CZ[47]. They observed this contraction between one and ten minutes after light onset and during continuous light exposure. That time scale is very different from ours and that of Tan et al. (2024), but suggests that there is a slow, late contraction of the CZ. In their second investigation, they used Gaussian mixture modeling to identify multiple contributions to each of the complex peaks observed in the axial reflectance profile of a commercial OCT image[48]. They used this model to monitor the movement of multiple reflective layers. Interestingly, they show that the ISOS is fit best by a mixture of three Gaussian functions, which they term ISOS-1G, ISOS-2G, and ISOS-3G, with the next peak outward attributed to ROST. Based on unpublished AO-OCT images acquired in our lab at a similar location, we feel confident that the relatively dim band termed ISOS-3G is due to the OS tips of the sparsely distributed cones, an attribution they independently adopted in subsequent work[49]. Treating ISOS-1G as our ISOS, ISOS-3G as our COST, and their RPE as our RPE, and digitizing and subtracting the movements of these bands relative to ELM, we were able to infer changes in OS, CZ, and SCS. These differences indicated a contraction of cone OS and CZ between 1 and 5 minutes after light onset, and elongation of cone OS and CZ between 5 and 10 minutes after light onset. The SCS appeared to contract between 1 and 7 minutes after light onset. In their third study, again using a commercial OCT system, they monitored slow light-evoked dynamics of the ELM-ROST and ELM-RPE distances, and computing the ROST-RPE distance, which they term the SRS[49], but which we have called the SCS. They showed reproducible contraction of the SCS between 1 and 10 minutes after light onset, with a magnitude of −20 %.

## 6 Conclusion

In this work we have shown that visible stimuli cause relative movement among multiple layers in the outer retina. The results above suggest that three compartments change in thickness after stimulation: the outer segment of the photoreceptor contracts briefly and then elongates; the supracone space, between the outer segment tip and the RPE, elongates briefly and then contracts; and the ciliary zone, between the ISOS and RPE, contracts briefly and the elongates, both with lower amplitude than the outer segment.

The first of these observations confirms previous reports of OS deformation after light stimuli. The latter two observations have not been reported before, but are consistent with light-dependent changes in the hydration of the extracellular matrix in the subretinal space. Because those hydration changes take place in a context of water movement through the retina, and because water clearance from the RPE is known to be impacted by disease, the ability to measure them noninvasively could be used as a novel assay of outer retinal health.

## Acknowledgments

We would like to thank: Susan Garcia (UC Davis) for her efforts in patient coordination; Profs. John S. Werner, Paul A. Sieving, Glenn Yiu, Ala Moshiri, Kareem Moussa (UC Davis) and colleagues Ratheesh Meleppat and Ankur Kumar (UC Davis) for helpful discussions about photoreceptor function, disease, and technical issues; Prof. Yifan Jian (OHSU) for development of OCT acquisition software.

## Data Availability Statement

The datasets generated and analyzed during the current study are not publicly available at this time but are available from the authors upon request.

